# Effects of focal epilepsy and aging on discrimination and reactivation of memories

**DOI:** 10.1101/2024.05.13.593984

**Authors:** Xin You Tai, Arjune Sen, Masud Husain, Sanjay Manohar

## Abstract

How are memories stored and subsequently retrieved? Pattern separation and pattern completion are distinct neurocognitive hippocampal processes that are thought to facilitate encoding and retrieval of memories. However, the way in which these operations affect the quality of memory remains unclear and there is currently no single behavioural task in humans to measure both mechanisms simultaneously. Here, we describe a novel paradigm which provides a distinct measure of each process and apply it to people with focal epilepsy, a condition associated with hippocampal dysfunction, comparing them with healthy young and older participants. Pattern separation and pattern completion were observed in healthy younger individuals. A specific deficit in pattern completion was identified in people with focal epilepsy, while the healthy older cohort showed degraded ability to perform pattern separation. To understand the underlying mechanisms, we simulated human performance using an auto-associative neural network. Modelling indicated that disruption in different populations of neurons could explain the observed memory profiles. These findings show that pattern separation and pattern completion are distinct processes that can be measured from a single behavioural task and are differentially affected by focal epilepsy and aging.

## Introduction

Pattern separation and pattern completion are two specific and complementary computational operations that facilitate storage and retrieval of information in our memory^1^. Pattern separation is the process whereby the neural representations of similar stimuli are individuated and kept separate, so each item can later be recalled separately^2^. Pattern completion is the process of restoring entire and accurate neural representations of such stimuli when presented with a partial or degraded cue. Both processes are considered to be crucial for our day-to-day memory^3^.

Theoretical and computational models describe distinct hippocampal circuitry implementing each of these operations, so that they can be performed independently^1,4–6^, and experimental animal models have provided some corroborative evidence^7,8^. However, a fundamental question that remains is how these two theoretically distinct processes map onto human behaviour. Current memory tasks tend to focus on one process or the other, making it difficult to disambiguate the two. Uncertainty partly arises because most paradigms have one behavioural outcome – wherein the participant is either correct or incorrect – but nevertheless attempt to extract two distinct measures from this performance.

**Pattern separation** is commonly operationalized using memory tasks where the test stimulus is similar – but not identical – to an originally remembered stimulus. A popular pattern separation paradigm, named the ‘Mnemonic Similarity Task’ (MST), is a change detection task in which participants identify whether the probe is the same (*repeat*), similar (*lure)* or different (*foil/new*) to the encoding set^3^. There may be different levels of subjective similarity between original objects and their lures. A ‘lure discrimination index’ is calculated based on accuracy of correctly identifying *lures*^9^ with most studies correcting for a bias in responding ‘similar’ by subtracting the rate that new objects are identified as ‘similar’^10,11^. Other approaches, however, calculate this pattern separation index from the *lure* accuracy in relation to false alarm rate (where similar items are reported as ‘old’)^12,13^ and one experiment accounted for errors in both (responding ‘old’ or ‘new’ to similar lures)^14^. This inconsistency is further compounded by some authors considering the false alarm rate as a metric for erroneous pattern completion activity, or ‘over-activity’^15–18^. However, lure accuracy used to calculate pattern separation is directly related to false alarm rate and, therefore, considering this as a measure of pattern completion is likely to be an over-interpretation of the behavioural response. There may, for example, be other reasons for this error including poor encoding of the original stimulus or forgetting due to intervening memory interference.

**Pattern completion,** by contrast, has been measured using memory recall cued by a degraded or masked stimulus. For example, the ‘Memory Image Completion’ (MIC) task uses a visual delayed recognition paradigm where participants learn a set of scenes and are later asked to identify the learned scenes, with new scenes mixed in, which are masked to different degrees^19^. Accuracy of identifying learned scenes decreases when more of the scenes are masked. A pattern completion metric termed as a ‘bias towards completion’ was derived by subtracting the accuracy of identifying a new scene (not present during the training phase) from the accuracy of identifying previously learned scenes. While this measure could arise from incorrectly completing a previously stored scene, it may also be confounded by incorrectly discriminating a degraded new scene and thus reflect both processes of pattern separation and completion. Therefore, current tasks in the literature do not cleanly isolate each cognitive process. Further, while pattern separation and pattern are typically studied in episodic memory, the timeframe for these processes is unclear and may also apply to short term memory. Accordingly, some aspects of WM engage the hippocampus, especially temporal context ^20,21^. Therefore, WM might maximally engage PC/PS mechanisms if information also needs to be retained in the longer term.

Nevertheless, pattern separation and pattern completion offer an intriguing framework for encoding and retrieval of information in health and disease^22^. Understanding how these key memory processes are perturbed in individuals with hippocampal damage may offer important insights into memory. Two conditions examined in this regard include aging, a natural process associated with a decline in hippocampal function and structure^23^, and Alzheimer’s disease, the most common neurodegenerative dementia whereby early pathology occurs in the hippocampus^24^. Studies in older individuals and people with Alzheimer’s disease report reduced pattern separation which has been attributed to ‘over-pattern completion’ ^11,15,25^. As explained above, this conclusion may not be fully justified because, in the mnemonic similarity task used, the measure of pattern separation is directly related to pattern completion.

Focal epilepsy, in which seizures begin in one part of the brain before spreading, is another important condition that may provide insights into pattern separation and pattern completion. In adults, focal seizures most commonly originate from the temporal lobe,^26^ known as temporal lobe epilepsy, but other non-temporal lobe focal epilepsies also affect abnormal networks involving the hippocampus^27–29^. Thus the hippocampus may be an important hub across focal epilepsies. A growing body of evidence now also suggests a bi-directional relationship between focal epilepsy and Alzheimer’s disease with histopathological^30,31^, imaging^32^, and electrographic intersections^33^. Thus, while Alzheimer’s disease increases the risk of developing focal seizures, the converse also holds: people with focal epilepsy are at greater risk of developing Alzheimer’s disease^34,35^.

A few behavioral studies in people with focal epilepsy have shown reduced pattern separation^36,37^, while one investigation of individuals with hippocampal damage using two sequential tasks of pattern separation and pattern completion described a deficit in both processes compared to controls^38^. However, using sequential tasks does not overcome the issue of pattern completion or pattern separation being inadequately controlled for in each task. A further issue is that most pattern separation and completion metrics are dependent on overall task accuracy, which may be affected by attention or executive function. These non-mnemonic cognitive factors may be generally lower in ageing and people with neurological conditions, such as epilepsy and dementia, and are harder to interpret in the context of healthy controls. Therefore, having a distinct measure of each operation may offer a useful test of hippocampal function.

One key attraction of studying pattern separation and completion is that they offer a computational framework that maps to neurobiology ^5,39–41^. Computational models of memory provide controlled environments for assessing the complex and interactive nature of information encoding and retrieval in the brain and also allow testing of predictions in a parameter space beyond what is feasible or easily achieved with human or animal participants. Importantly, computational models can move outside of physiology to simulate disease and “the patient”. Equipping recurrent artificial neural networks with associative plasticity gives a content-addressable memory system^42^ that can emulate the functional properties of human memory ^43^. This can be used to generate neural and behavioural hypotheses about pattern separation and completion.

Here, we describe a novel ‘Occlusion at Retrieval Task’ (OART) which simultaneously assesses both pattern separation and pattern completion within the same paradigm. Our metrics of each process correct for differences in overall memory accuracy, with both derived from retrieving information about different dimensions of the *same* probe. The task was tested in a cohort of healthy young and older participants, and individuals with focal epilepsy to identify whether overall memory performance can be explained by specific deficits in either of these cognitive processes. Further, we apply computational modelling of our behavioural results using a modified auto-associative Hopfield network^44^ to better understand potential underlying mechanisms of memory encoding and retrieval.

## Results

### Novel cognitive task with distinct measures of pattern separation and pattern completion

In the OART each short term memory trial began with an encoding phase, where participants were presented with a shape to remember (**Figure 1a**). The shape had four arms which could be either long or short. After a delay, there was a first retrieval phase, where they saw a probe which was either the same as the stimulus at encoding (50% of trials), similar with one arm different (more similar, 25%) or two arms different (less similar, 25%, **Figure 1b**). The probe was either more covered (two arms occluded) or less covered (one arm obscured). Participants were asked whether the probe was either the *same* or *different*. This design provided distinct assessment of **pattern separation**, based on two similarity levels, and **passive pattern completion** based on two levels of probe occlusion. Crucially, these two manipulations (similarity and occlusion) were performed on different dimensions of the same stimulus.

**Figure 1.**
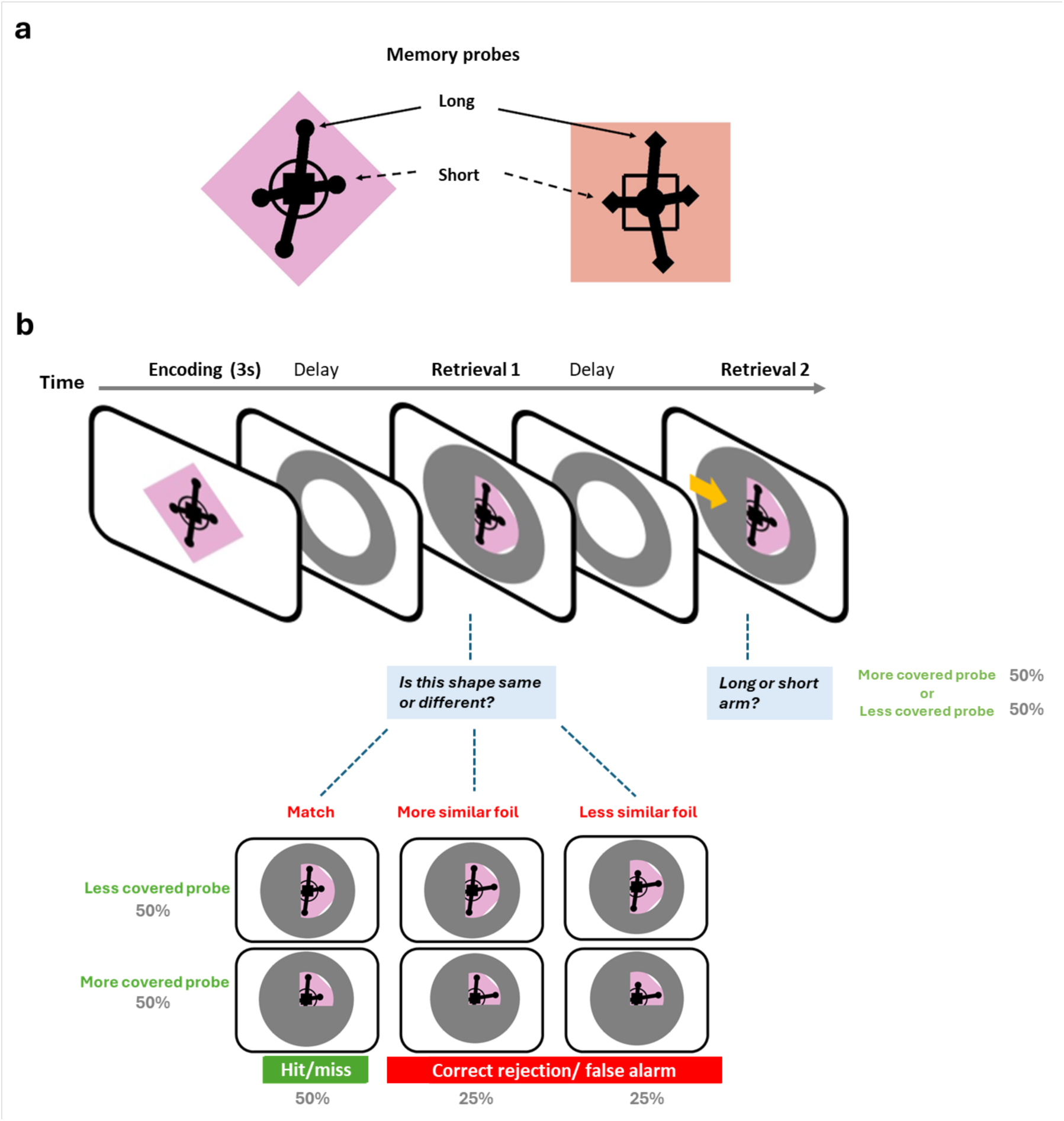
Occlusion at Retrieval Task design. **a** Example of two stimuli used in the Occlusion at Retrieval Task. Each stimuli had four arms which were either long or short. These were used to manipulate similarity between encoding stimuli and probe. Other features included a coloured background shape as well as a middle, intermediate and end-arm shape which were not changed within a trial. **b** Task design showing a single shape to-be-encoded followed by a first probe at Retrieval 1. This would either be an exact match, more similar foil (different by one arm) or less similar foil (different by two arms). Different levels of similarity assess **pattern separation**. Participants were asked whether this shape was the same or different to the shape shown at encoding. Probes were either more or less occluded/ covered to assess **passive pattern completion**. This was followed by a second delay before the same probe with an arrow over a covered arm was presented at Retrieval 2. Participants were asked whether the arm that the arrow pointed to was long or short, in order to probe **active pattern completion**.

In a second retrieval phase, participants saw the same probe but this time were asked to indicate whether a covered arm was *long* or *short,* as a measure of **active pattern completion** (**Figure 1b**). Note that two probes were deployed at retrieval to assess how well pattern completion occurred passively (as when an experience triggers a recollection of related things), or actively (as when we consciously try to remember something based on a cue).

Models of hippocampal encoding allow one-shot learning^45^. Therefore, in this study, we focused on memory over short durations. Longer-term memory probes were also included (every fourth trial) to assess episodic function on this task (**Figure S1).** Here, the stimulus from four trials ago was probed. The colour and shapes provided a context indicating which trial was being probed. The probe was either identical to the encoded stimulus or had all four arms of different length to maximise discriminability.

The OART was first tested in a healthy young cohort (N=48, mean age 27 years, range 19-34) to establish baseline performance. Following signal detection theory, this control group demonstrated a high ability to discriminate between same versus different probes (d’ = 3.58) as well as low bias (c=0.05, not significantly different to zero) across all trials. For non-match trials, as expected, accuracy for less similar probes was greater than more similar probes [F(1, 94) = 11.30, p=0.001] which represented pattern separation (**Figure 2a**). For match trials, accuracy was higher for probes which were less occluded [t=-3.15, p=0.028], indexing passive pattern completion (**Figure 2b**). A small difference in accuracy between more similar and less similar conditions indicates better pattern separation, so this measure is a relatively conservative index. A small difference between more covered and less covered probes indicates better pattern completion, i.e., that covering an arm is less detrimental to performance. Again, this represents a relatively conservative measure.

**Figure 2.**
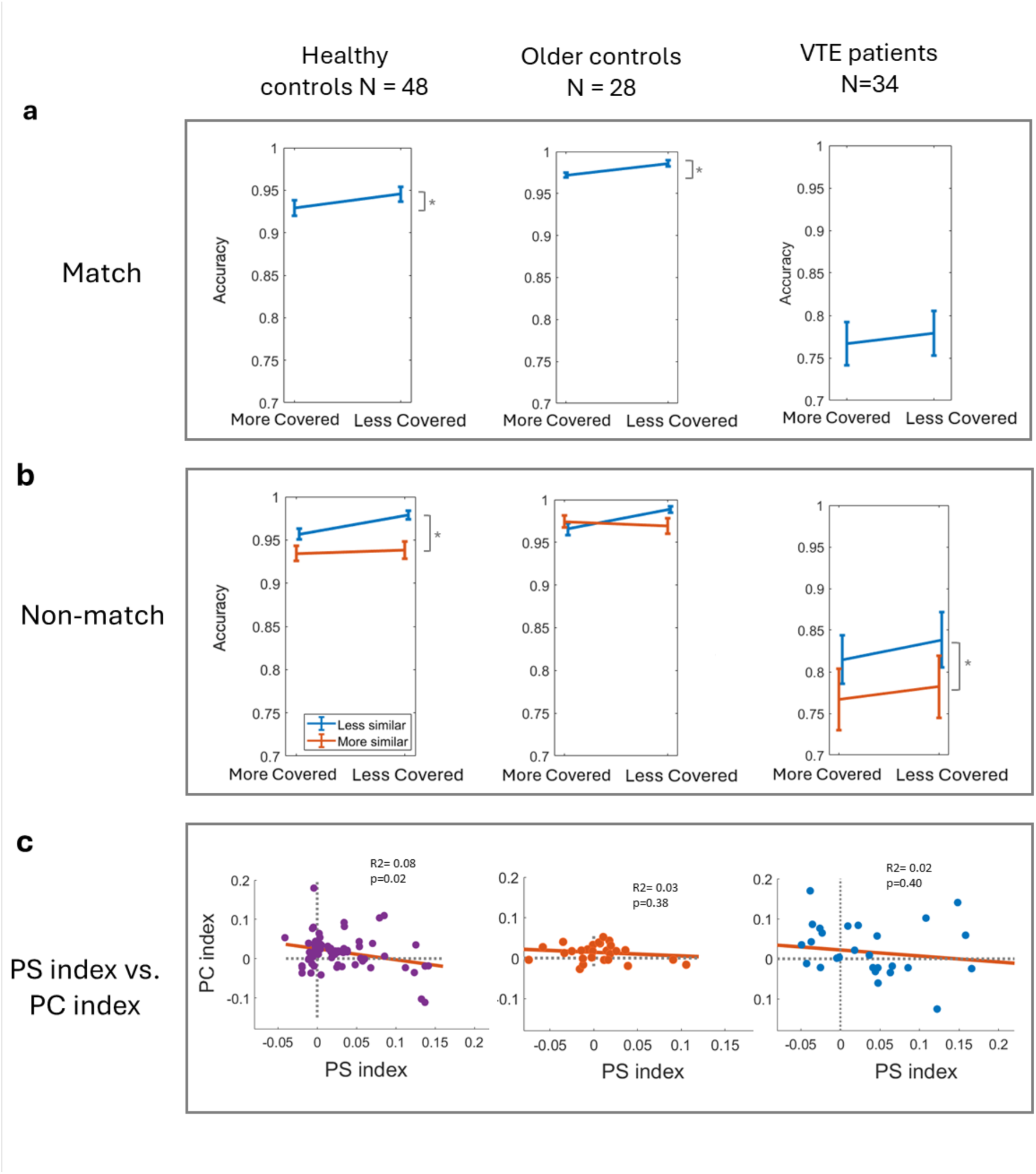
Occlusion at Retrieval Task results across healthy young, older and people with focal epilepsy. **a** Accuracy was significantly higher in match trials for probes which were less covered, reflecting pattern completion, in the healthy young (p=0.001) and healthy older cohorts (p=0.012) but not for people with focal epilepsy (p=0.078). **b** For non-match trials, healthy young (p=0.02) and people with focal epilepsy (p=0.042) had higher accuracy for probes which were less similar, reflecting pattern separation. **c** Correlation between PS index and PC index across the three cohorts showing a weak correlation for healthy young individuals only (r^2^= 0.08, p = 0.02, PC and PS index points have been jittered by 2% to help distinguish clusters). *PS- pattern separation, PC- pattern completion*.

There was a weak negative correlation between these pattern separation and pattern completion indices (r^2^= 0.08, p = 0.02), calculated by the difference in accuracy based on probe similarity and probe completeness respectively. This indicated that the pattern separation and pattern completion indices were two distinct but related metrics (**Figure 2c**). Analysis of the active arm completion probe showed a higher accuracy with stimuli which were less covered (t = 2.17, p = 0.026), also consistent with a pattern completion process (**Figure S2a**).

### Dissociable effects of aging and focal epilepsy on pattern separation and pattern completion

The OART was then tested in healthy older individuals (N=28, mean age 65 years, range 45-77) and people with focal epilepsy (N=34, mean age 34 years, range 19-59). The age distribution for the focal epilepsy cohort spanned across the age range of both younger and healthy controls. We further control for age as a covariate in our main model. Across all trials, while both healthy younger and older individuals’ performance was associated with *d*’ values of >4 and bias <0.05 (not significantly different from zero), people with epilepsy had a *d*’=2.44 and bias = -0.13 (significantly different from zero, t=-3.12, p = 0.028), the latter indicating a bias towards reporting a stimulus was the same. Accordingly, the epilepsy cohort had a significantly greater mean accuracy on match trials compared with non-match trials (accuracy of 0.87 vs 0.80 respectively, t=2.87, p = 0.007 when testing the difference in means).

To assess overall performance across groups, we used an *a priori* designed trial-level generalised logistic mixed effect model, predicting accuracy from match (vs. non-match) trials, probe similarity (less vs. more), probe cover (less vs. more), epilepsy patient group (vs. controls), age and sex, plus interaction terms of patient group*match trials, patient group*probe cover and patient group*probe similarity (**Figure S3**). Main effects demonstrated significantly higher accuracy with match trials (t=3.13, p=0.002), probes being less covered (t= -4.68, p<0.001), probes being less similar (t= 5.82, p<0.001), and being a healthy control (t= -6.08, p<0.001) and of older age (t= 2.31, p=0.02). The model also identified a significant interaction between match trials and being in the epilepsy group (t= -6.83, p<0.001), consistent with people with focal epilepsy having a bias to choose ‘same’ that was noted above. There was also a significant interaction between probes being less covered and being in a control group leading to higher accuracy (t= 2.20, p=0.023) which reflects that healthy individuals were better when the probe was less occluded, but this was not observed in people with focal epilepsy. There was no significant interaction between probe similarity level and being a person with focal epilepsy (t= -0.88, p=0.375).

To understand the interactions further, two-factor ANOVA was performed for each cohort on match and non-match trials with main effects of similarity level and probe completeness. Performance on match trials (**Figure 2a**) was more accurate for less covered probes compared to more covered probes and the effect was statistically significant in healthy young [F(1, 94) = 11.30, p=0.001] and healthy older individuals [F(1, 54) = 6.25, p=0.012] but not in people with epilepsy [F(1, 64) = 2.58, p=0.078]. This may reflect a pattern completion effect in healthy young and healthy older individuals. For non-match trials (**Figure 2b**), accuracy was higher when probes were less similar to the original, compared to more similar, in healthy younger [F(1, 94) = 5.43, p=0.02] and people with focal epilepsy [F(1, 64) = 4.83, p=0.042] but this was not the case in healthy older individuals [F(1, 54) = 0.45, p=0.50]. This may reflect a pattern separation effect in healthy young and individuals with focal epilepsy.

Next, we asked whether separation and completion were independent. Pattern separation index was weakly correlated with pattern completion index for healthy young individuals, as noted previously, but not for healthy older individuals or people with focal epilepsy (**Figure 2c**). The Bayes Factor suggested weak evidence towards the null hypothesis for each of the latter two cohorts (BF01 = 5.36 and BF01 = 4.87, respectively). To account for possible ceiling effects, an arcsine transform was applied to the data but this did not change the associations between pattern separation index and pattern completion index in any of the cohort groups.

### Temporal lobe epilepsy showed similar deficits to all participants with focal epilepsy

The generalised logistic mixed-effects model was run with only people with temporal lobe epilepsy (N=28), excluding those with frontal lobe epilepsy (N=5) and focal epilepsy of unclear origin (N=1). Model outputs were similar with a significant interaction between match trials and being in the epilepsy group (t= -6.18, p<0.001) and a significant interaction between probes being less covered and being in a control group leading to higher accuracy (t= 1.98, p=0.042). The main effects also demonstrated higher accuracy with match trials (t=2.59, p=0.009), probes being less covered (t= - 4.68, p<0.001), probes being less similar (t= 5.94, p<0.001), being a healthy control (t= -6.54, p<0.001) and of older age (t= 2.45, p=0.02). Therefore, in this model, having temporal lobe epilepsy had a similar negative effect on memory performance compared to when all individuals with focal epilepsy were included into the model (**Figure S4**).

### Active completion, longer term memory probes and reaction time

In the healthy older cohort, accuracy was higher in the active arm completion probe when stimuli were less covered (t = 2.35, p = 0.026), consistent with a pattern completion process described above. For the focal epilepsy cohort, there was no significant difference between active arm completion accuracy, for more or less covered probes (t = 1.33, p = 0.192).

Performance was uniformly poor for longer term memory probes. In the healthy younger cohort, overall accuracy for longer-term probes was 0.62 (SE = 0.05) with five individuals scoring below chance level (0.5) (**Figure S2b**). The healthy older cohort had an overall accuracy of 0.61 (SE = 0.02) with one individual scoring below chance level (0.5) while the focal epilepsy cohort had an overall accuracy 0.54 (SE = 0.02) with six individual scoring below chance level (0.5). Despite the discrimination being easier, performance with longer-term memory probes was significantly worse than short term memory probes (p <0.001 in all groups), indicating that information was lost from memory over four trials.

Average time taken to give a response for match and non-match trials were similar within each cohort group (no significant difference) as younger individuals had the fastest average reaction time while people with focal epilepsy took the longest to respond (**Figure S5**). When plotting reaction time against accuracy at a trial level for each cohort, a speed-accuracy trade-off was observed with the healthy older cohort demonstrating a longer reaction time corresponding to higher accuracy (**Figure S5**). Importantly there were no RT differences between the level of similarity or between the level of completeness (p>0.05).

### Modified Hopfield network modelled responses to pattern similarity and probe completeness

We used an auto-associative, also known as a Hopfield, network to encode and retrieve the features present in our stimuli, encoded as a binary vector^45^. Stimuli were encoded by activating the pattern of neurons corresponding to the features present in the visual stimulus, then updating the weights between the neurons according to a Hebbian rule to store the pattern auto-associatively. Visual features were either ’task’ neurons for features to be reported, i.e. corresponding to arm length in the behavioural task, or ‘context’ neurons, corresponding to irrelevant features which differ for each new stimulus such as colour or central shape. The response of the network to a probe was calculated by activating the neurons corresponding to the features of the probe that were uncovered and reading out the resulting activation pattern. We derived a *model certainty* measure indicating how strongly the network recognised the retrieval probe to be the same as a previously stored stimulus (**Figure 3a**). This was based on the strength with which each individual neuron/ pattern component was active or inactive (**Figure S6**).

**Figure 3.**
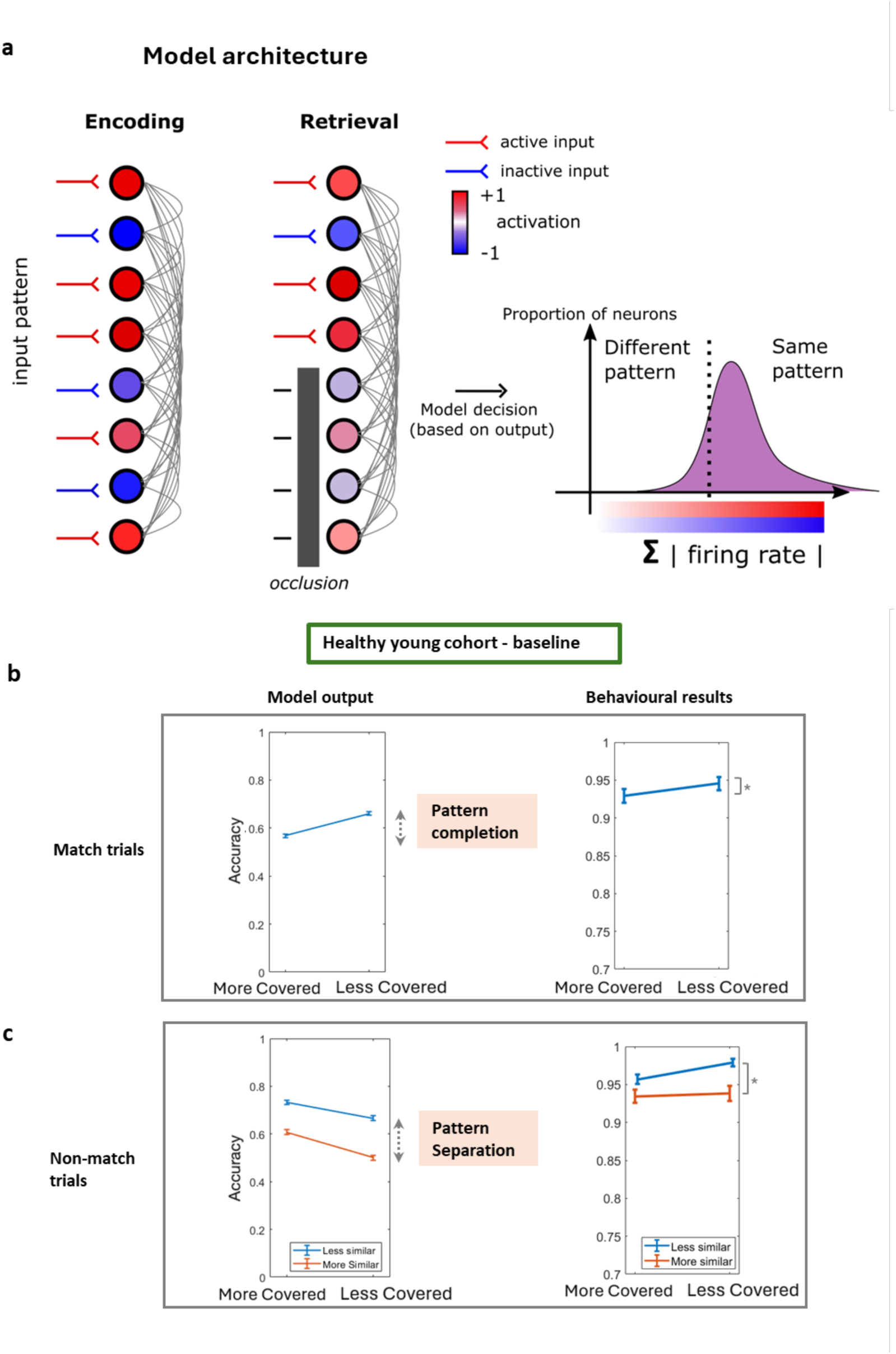
Modelling the Occlusion at Retrieval Task in “healthy young” individuals. **a** Model architecture showing an input pattern at encoding stored within synaptic weights/ firing rate of the fully connected recurrent neural network. During retrieval, a partially covered pattern is presented to the network. A model decision is given based on the sum of the absolute firing rate to output either “same” or “different”. **b** Match trial model performance shows higher accuracy for less covered probes compared to more covered probes (pattern completion effect), consistent with behavioural results from the Occlusion at Retrieval Task. **c** For non-match trials, the model performed better for less similar probes reflecting a pattern separation effect also seen in healthy younger individuals.

Closely matched experimental parameters were used to model the Occlusion at Retrieval Task. Specifically, 196 stimuli were encoded with corresponding match probes, more similar non-match probes (25% of the pattern flipped) and less similar non-match probes (50% flipped). All retrieval probes were either more covered (50%) or less covered (25%) and conditions were counterbalanced. The key output was whether the model considered the probe as being ‘same’ or ‘different’ compared to encoding stimuli. The threshold for choosing “same” or “different” for a probe set as the median firing rate reflecting the assumption that participants would generally choose 50% same and 50% different in a standard behavioural task. As a baseline model architecture, we used N = 100 neurons with an equal number of context neurons. The model has no other free parameters.

The model output showed worse performance in match trials for more covered probes and was therefore more ‘certain’ of a probe being a match when more information was present. This output matched the behavioural results on the OART and represented pattern completion (**Figure 3b**). For non-match trials, model performance was better for probes that were less similar to the encoding stimulus, which also matched behavioural results and represented pattern separation (**Figure 3c**). Interestingly, the model predicted better performance at choosing “different” for non-match probes when they were more covered, which may reflect a lower likelihood of completing the pattern accurately when the probe is more obscured.

### Fewer context neurons model healthy older group while fewer task neurons simulated focal epilepsy group

Reducing the number of context neurons was the model parameter manipulation that best modelled the behavioural results of the healthy older cohort in the Occlusion at Retrieval Task. With half the context neurons available, the model still showed better performance in match trials for less covered probes compared to more covered probes (pattern completion effect). This was observed in the behavioural results (**Figure 4a**). For non-match trials, however, there was no clear difference in performance for more or less similar probes suggesting no pattern separation effect. This also corresponded with the behavioural results (**Figure 4b**) suggesting different underlying biological computations with aging.

**Figure 3.**
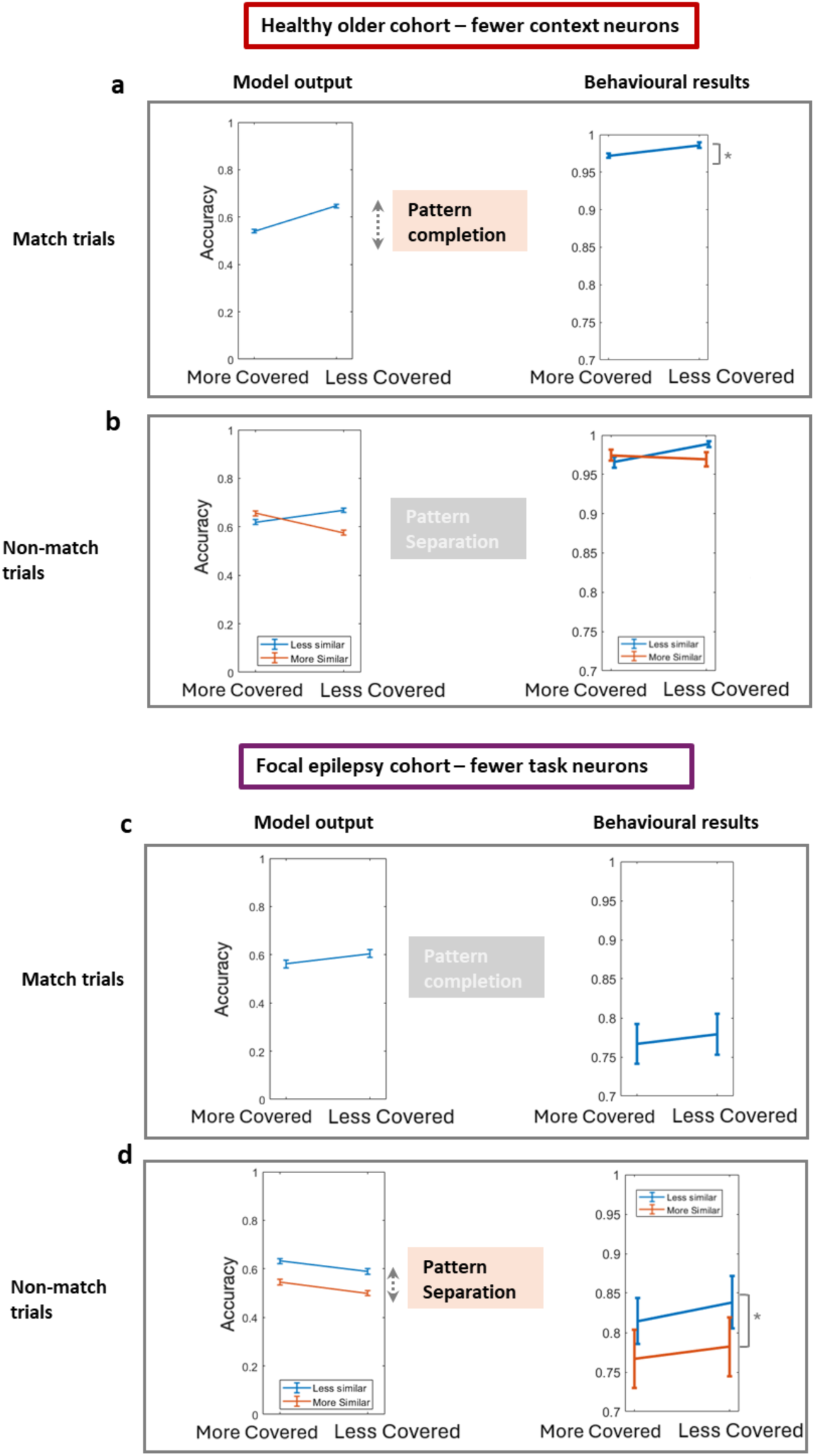
Modelling Occlusion at Retrieval Task (OART) in healthy older and focal epilepsy cohort. The healthy older cohort behaviour was simulated by, **a**, reducing the number of context neurons led to higher match trial model performance for less covered probes compared to more covered probes (pattern completion effect). **b** For non-match trials, different levels of probe similarity were not associated with an overall performance difference or clear pattern separation effect. Results from the focal epilepsy cohort was best simulated by reducing the number of task neurons which, **c**, resulted in a smaller pattern completion effect (when performance is better with less covered probes). Correspondingly, behavioural results did not show a significant difference in accuracy between more and less covered match probes. **d** For non-match trials, better performance was associated with less similar compared to more similar probes.

The behavioural results of people with focal epilepsy were simulated by lesioning the pool of task neurons. When considering a model with half the number of task neurons, the model maintained better performance for non-match trials with less similar probes representing a pattern separation effect that was observed in people with focal epilepsy (**Figure 4d**). For match trials, however, the effect of probe cover was smaller with reduced task neurons as the model was only slightly better at identifying less covered match probes (**Figure 4c**). People with focal epilepsy had a trend towards better performance with less covered match probes however this was not statistically significant. A larger sample size may be needed to detect this effect which was predicted by the model.

The simulations outlined above demonstrate that the auto-associative network can simulate the Occlusion at Retrieval Task and that the dissociable results from healthy older people and people with focal epilepsy may be accounted for by lesioning different pools of neurons within the network. This may reflect underlying biological mechanisms in both cohorts.

## Discussion

The Occlusion at Retrieval Task (OART) provides a novel approach to obtaining distinct measures of pattern separation and pattern completion from a single memory paradigm by manipulating different dimensions of the same stimuli in a single task. Healthy younger individuals demonstrated that probes with higher similarity and lower completeness were associated with worse performance, reflecting pattern separation and completion processes, respectively. These indices had a weak negative association which rebuts the idea that they lie on opposite ends of a unitary spectrum^15^.

Examination of these metrics in older individuals and people with focal epilepsy provide specific insights on pattern separation and pattern completion. While previous studies suggest that older individuals perform less well in change detection tasks due to ‘under-separation’ and ‘over-completion’ ^15–18^, results from the present experiment indicate a similar level of pattern completion to that seen in healthy younger individuals, but that less pattern separation occurred. The better average performance of the healthy older cohort compared to younger cohort in our paradigm may have been due to a speed-accuracy trade-off. This emphasises the need for pattern separation and pattern completion metrics that are not dependent on accuracy.

People with focal epilepsy, including only those with temporal lobe epilepsy, performed worse overall compared with healthy individuals but still showed better discrimination of probes which were less similar, reflecting a pattern separation process. By contrast, level of probe cover did not have a significant effect on performance which may reflect a deficit in pattern completion. This is the first report of a dissociable deficit between these two processes in older healthy individuals and those with focal epilepsy. These results were unchanged when considering those with temporal lobe epilepsy only which is associated with hippocampal dysfunction. As such, the current work supports previous findings that temporal lobe is often involved in distributed seizure networks, even when abnormal electrical activity begins elsewhere in the brain^27–29^.

The OART offers more specific and matched measures of pattern separation and pattern completion compared to previous tasks, and does not solely relate to overall memory accuracy. This is important as patients with neurological conditions may often perform worse than healthy controls which, in current tasks described in the literature, would usually be erroneously observed as deficits in pattern separation or pattern completion. Therefore, the OART improves upon several current tasks in the literature.

A modified Hopfield, or auto-associative, network was able to simulate results of the OART, which characterizes the separable computations underlying pattern separation and pattern completion. Compared to other autoassociative models of memory, we derived a same-different decision directly from the neurons in the network. Better performance identifying “same” probes was observed when match probes were less covered (pattern completion effect) while, in non-match trials, the model performed better in choosing “different” with less similar probes. Changing different model parameters led to outputs that simulated behaviour from healthy older participants (decreasing context neurons) and people with focal epilepsy (decreasing task neurons). This corresponded with reduced pattern completion in healthy older participants and reduced pattern separation effect in people with focal epilepsy.

Our finding supports the view that pattern separation and completion are distinct processes in which different neuronal populations are involved. Remarkably, this model performed these two separate processes using a single neuronal population. This may be surprising when considering that, neuroanatomically, the dentate gyrus is hypothesized to perform pattern separation whilst CA3 is considered the site of pattern completion^45^. Our findings are, though, consistent with the hippocampus performing multiple independent processes as a single unit and helps resolve the uncertainty of how neuronal signals move neuroanatomically from entorhinal cortex through dentate gyrus to CA3 for pattern completion to occur without pattern separation or if there must be an additional perforant pathway between entorhinal cortex directly to CA3 to facilitate pattern completion without engaging dentate gyrus neurons.^46^

Interpreting the role, or biological relevance, of each neuronal pool is not straightforward. Task neurons are directly involved with retrieving stored representations within the model. Reducing or lesioning task neurons may be analogous to disrupting the memory retrieval process which may occur in the CA3 region of the hippocampus. CA3 has numerous recurrent synaptic collaterals ^47^ and is vulnerable to epileptic aberrant activity. Task neurons are, however, also required to encode stimuli in the first instance. Therefore, both encoding and retrieval are likely affected in people with focal epilepsy, although a reduced pattern completion effect was more clearly observed in both behavioural results and predicted by the model output.

Changing context neurons modelled the behaviour of older participants. While not directly involved in the relevant task pattern, context neurons allow better encoding of each pattern by increasing the difference between other patterns in the model. This, in effect, describes a pattern separation effect which was reduced in the behavioural results of older participants. From a computational perspective, the context neurons allow patterns to be stored separately within more distinct basins of attraction within the network^8,48^ reducing the chance that one pattern will be confused by another pattern at recall. This change in context neurons may reflect generalised atrophy observed in the ageing brain^49^.

It is important to consider limitations to the current behavioural paradigm. While the stimuli were grounded in a mathematical framework that was designed to precisely manipulate similarity and probe completeness, it could be argued that their abstract nature will make it difficult to employ in a more episodic paradigm with longer delay and more interference between encoding and retrieval. Adding more context and complexity to the probes will help with this. Also, collecting baseline cognitive measures in healthy participants would have been preferred (but this was not possible owing to disruption to in-person testing as the study period coincided with the Covid-19 pandemic).

In conclusion, we demonstrated a novel paradigm to assess both pattern separation and pattern completion within a single task. Results suggested that these two cognitive processes do not lie on a unitary spectrum, but are two related yet distinct operations. This is consistent with theoretical models and helps to resolve an existing debate in the literature. Pattern separation and pattern completion may be a useful framework of memory encoding and retrieval to help to understand changes in health and disease with dissociable effects of age and focal epilepsy.

### Methods Ethics

Ethical approval was granted by the University of Oxford ethics committee (IRAS ID: 248379, Ethics Approval Reference: 18/SC/0448). All research was performed in accordance with university guidelines, including participant research performed in accordance to the Declaration of Helsinki. Participants gave written informed consent prior to the start of the study.

### Participants

There were three participant cohorts tested in this experiment. A group of healthy young individuals with forty-eight participants (34 female, mean age 27 years, range 19-34) were recruited via a local database or the online platform Prolific. Participants were paid £10-15 for the task which was performed on a desktop computer under supervision or via the online platform Pavlovia. A healthy older group of 28 volunteers (18 female, mean age 65 years, range 45-77) were recruited via a local database and the task was done at home via the online platform Pavlovia. Three older volunteers did not understand the task properly and were excluded from the study.

A group of 38 people with focal epilepsy (20 female, mean age 34 years, range 19-59) were recruited from the epilepsy video telemetry unit, John Radcliffe hospital, Oxford, United Kingdom. In terms of seizure semiology, 28 individuals had temporal lobe epilepsy, five individuals had frontal lobe epilepsy while one individual had focal seizures of unclear origin. These participants were admitted to hospital for at least one week to have continuous monitoring of seizure activity. Most of this group had their anti-epileptic medications reduced during their hospital stay with the majority being tested off medication. Further demographics can be found in **Table S1,** including seizure localisation and underlying pathology. Four people with epilepsy were excluded as they did not complete the task due to fatigue or a seizure occurring during the task. Addenbrooke Cognitive Examination (ACE-III) was performed in the epilepsy group but not the other cohorts due partly to logistical reasons as face-to-face testing was not carried out for a substantial proportion of the study due to Covid-19 lockdown protocol.

### Occlusion at Retrieval Task

The ‘Occlusion at Retrieval Task’ was a novel pattern separation and pattern completion short term memory paradigm which manipulated similarity between encoding stimuli and retrieval probe in addition to covering one or two dimensions of the probe. Memory stimuli followed a common base shape with four dimensions/ ‘arms’, either long or short (fixed lengths of 140 or 60 pixels), with a random angle of 15 degrees to either side of the base arm (**Figure 1a**). The crucial manipulation of similarity was achieved by altering arm lengths between encoding stimuli and retrieval probe on each trial. To create complex and visually rich stimuli, additional features were added: a coloured background shape (eight colours, four possible shapes– square, circle, diamond and hexagon) as well as end-arm, central and middle shapes (four possible shapes for each). These contextual features were held constant throughout one short term memory trial.

The Occlusion at Retrieval Task had three different probe stages:

1. **Short term memory change detection (main probe)** During each trial, participants would see one stimulus at encoding and, after a delay, would be presented with a single probe which was either the same as the stimulus at encoding, similar with one arm different (more similar) or two arms different (less similar). The probe was either more covered (two arms obscured) or less covered (one arm obscured). Participants were asked whether the probe was either the same or different (**Figure 1b**). This probe stage was designed to assess pattern separation (two different similarity levels) and passive pattern completion (two different levels of probe cover). This short term memory probe is the key condition described in this report. Each participant performed 196 trials of this main short term memory change detection probe with counterbalanced conditions of more or less similar, and more or less covered.
2. **Short term memory arm completion** Following their first response, participants were asked to indicate whether one of the covered-up arms was either long or short as a test of active completion.
3. **Longer-term memory** Finally, every four trials, a longer-term memory probe was shown and participants were asked if this stimulus was the same or different to a few trials ago. The full probe was shown (no arms were covered) and the arm lengths were either the all the same or all different. This was the largest contrast possible between stimuli and long term memory probe due to the difficult nature of this probe.

### Analysis

Group accuracy was the dependent variable for rm-ANOVA or with a Student’s T-test to compare means between two groups. Signal detection theory was applied to calculate discriminability and bias. When appropriate, Bayes factor was calculated to examine for evidence towards the null hypothesis using an unbiased prior distribution, such as theta = beta(1,1) or a stretched version, using JASP (Version 0.16.3).

A ‘*pattern separation index*’ (PS index) was derived by taking the difference between mean accuracy of trials with less similar probes minus trials with more similar probes:

𝑷𝑺 𝒊𝒏𝒅𝒆𝒙 = 𝑨𝒄𝒄𝒖𝒓𝒂𝒄𝒚 𝒐𝒇 𝒍𝒆𝒔𝒔 𝒔𝒊𝒎𝒊𝒍𝒂𝒓 𝒑𝒓𝒐𝒃𝒆𝒔 − 𝒂𝒄𝒄𝒖𝒓𝒂𝒄𝒚 𝒐𝒇 𝒎𝒐𝒓𝒆 𝒔𝒊𝒎𝒊𝒍𝒂𝒓 𝒑𝒓𝒐𝒃𝒆𝒔

A ‘*pattern completion index*’ (PC index) was derived by taking the difference in mean accuracy between trials with less covered probes minus trials with more covered probes on trials where the probe was the same as the encoding stimuli:

𝑷𝑪 𝒊𝒏𝒅𝒆𝒙 = 𝑨𝒄𝒄𝒖𝒓𝒂𝒄𝒚 𝒐𝒇 𝒍𝒆𝒔𝒔 𝒄𝒐𝒗𝒆𝒓𝒆𝒅 𝒑𝒓𝒐𝒃𝒆𝒔 − 𝒂𝒄𝒄𝒖𝒓𝒂𝒄𝒚 𝒐𝒇 𝒎𝒐𝒓𝒆 𝒄𝒐𝒗𝒆𝒓𝒆𝒅 𝒑𝒓𝒐𝒃𝒆𝒔

The advantage of using performance differences, based on similarity and probe cover, was to a measure of pattern separation and completion regardless of overall memory performance.

Additionally, a generalised logistic mixed-effect model analysed trial—level performance to specifically investigate higher-level interactions within the experiment. The model was specified based on prior hypothesis using the fitglme function within MATLAB, Mathworks. Match (vs non-match trials), similarity level, cover level, patient group (epilepsy vs. healthy), age and sex were fixed effects while a subject ID was entered as a random effect. An interaction term was included between patient variable and match, similarity and probe cover level. Reaction time was investigated separately as a factor that may explain overall performance and also whether non-match trials were slower.

### Hopfield network model for memory encoding and retrieval

A Hopfield network of information encoding and retrieval has been previously used to model pattern completion that may occur in the hippocampus^50^. Memory representations are stored as orthogonal, or largely uncorrelated, patterns to increase network capacity^51^. For this study, we build upon previously described network architecture and propose modification to incorporate task and context neurons, interpretation of non-binary neuron activity and a novel, unbiased model decision on “same” and “different” stimuli.

Mathematically, an external input e_i_ is applied to each neuron i of the network through unmodifiable synapses to produce output firing y_i_. The output from each neuron is connected via recurrent collateral connections to the other neurons in the network and stored in a modifiable weights matrix wi_j_. This organization allows the initial activity of these neurons to be associated and ‘learned’ to model information encoding.

A modified Hebbian local learning rule is applied to the recurrent synapses in the network:

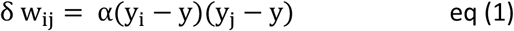

Where α is a constant and y approximates the average value of y_i_ and y_j_.

Subtracting the value y will prevent increasingly positive or negative, i.e. exploding, weight values.

Once this stimuli information is stored, a subsequent recall probe can be presented to the network as a full or partial cue as an external input to produce neural output firing in the pattern of the original ‘learned’ input.

During recall, an external input e_i_ activates neurons, and this activity propagates through the recurrent collateral system to produce an activation output of each neuron. The activation vector h_i_ is the sum of activations produced in proportion to the firing rate of each axon y_i_ operating through each modified synapse wi_j_.

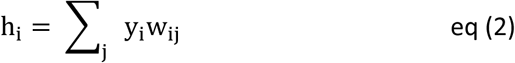

Where the sum is over the C input axons to each neuron, indexed by j. The output firing y_i_ is a function of the activation produced by the recurrent collateral activity (internal recall) and the external input (ei), where f is a non-linear (e.g. arc tan) activation function.

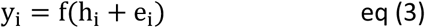

#### Model characteristics

### External inputs to the model

Following the classic Hopfield network literature, external input values fed into the network were generally constrained to either ‘1’ or ‘0’. This often depicts an ‘active’ or ‘inactive’ neuron and could also represent ‘long’ or ‘short’ arms of our visual stimuli to be stored into memory. We extend this to non-binary values to reflect a specific experimental parameter. For example, an external input of ‘0.5’ could represent a covered or uncertain arm length in the context of other inputs being either ‘1’ or ‘0’.

### Model sparseness

Sparseness level of model inputs is an important characteristic of an autoassociative network and has implications on the network storage capacity^44^. At a population level, sparseness S refers to the number of neurons which are firing at a given time. Therefore, S will range from 1/N to the S = 1 when all the neurons are firing, or ‘active’, and S = 0.5 if half the neurons are ‘active’ for a given neuronal pattern. To model the behavioural task, sparseness was fixed at S = 0.5 for this study.

### Task and context neurons

This model had two main neuronal populations. ‘Task’ neurons which were directly involved with encoding a pattern into the network and retrieving the stored representation. For example, the activity of one neuron may correspond to one component or feature dimension of the stimulus. By contrast, ‘context’ neurons were a second neuron population with random activations for each stimulus, presented together with the pattern to be remembered (with sparse level S=0.5). Despite not being directly involved in encoding or retrieving a specific pattern, context neurons helped to provide a stable representation of each pattern within the network. Conceptually, they can be considered as providing some related information such as an environmental association or potential spatial or temporal context specific to that pattern. Binding of remembered information into a unique environmental context, such as time, place and person, are often considered an essential attribute of hippocampal memories^52^. For this study, a baseline model architecture comprised of 100 task neurons (N=100) with an equal number of context neurons unless otherwise specified, for example, when the model was specifically ‘lesioned’ to simulate pathology.

### Modelling similarity

Pattern similarity can be manipulated in different ways. Following Euclidean geometry, the magnitude of the dot product between two vectors is a measurement of similarity whereby a higher value indicates greater similarity (this can also be represented by the cosine similarity) ^53^. To manipulate similarity between two patterns, we start with one pattern and took the inverse state of one neuron component (dimension) of the pattern, which we refer to as ‘flipping’ the component. For example, two similar patterns can be created by taking pattern A and flipping one component neuron state from ‘active’ to ‘inactive’ to create pattern B and leaving all other components the same. **Figure S7a** shows how pattern A is changed to pattern B (more similar) by flipping one active and one inactive neuron or to pattern B’ (less similar) by flipping all neuron states. As more components are flipped to create a new pattern B, the dot product between the patterns A and B decreases as they become less similar to each other. Generally, for the experiments outlined in this paper, a pair of inactive and active neurons were flipped at the same time to maintain the same model sparseness (S=0.5).

Similarity within a pattern can manipulated by repeating a series of component neurons within the pattern. For example, in **Figure S7b**, pattern A is comprised of 12 components with the first six components being the same as the second six components reflecting a high within-pattern similarity. Pattern A’ has a lower within-pattern similarity as the second six components are randomly different from the first six components.

### Model output: accuracy

The model output to a recall probe is a set of neural patterns that most closely matches the original input, based on stored model weights. These output patterns are generated from the firing rate of each neuron once the external input probe has passed through the recurrent collateral system. The representational space for the neuronal firing rate is continuous and will generally follow a gaussian distribution.

To convert the firing rates into binary patterns, the neuronal activity can be interpreted as an ‘active’ or ‘inactive’ neuronal state by applying a threshold activation value. Previous models will often use a threshold based on the sparseness value of the original stimuli^1^. For example, if original patterns had sparseness S = 0.5, then the threshold for being considered active was the median activation value of all neurons. Once each output neuron activity was interpreted as a 0 or 1, then this output pattern can be directly compared with the original learned patterns to calculate a ***model accuracy*** for a given set of probes i.e. how closely does the model output neural patterns match the learned input patterns for a given set of probes.

Our modified Hopfield network demonstrated several fundamental principles of memory encoding and retrieval including set size effect and retrieval probe completeness affecting model accuracy or other performance metrics (**Box 1**). Furthermore, it can also simulate results from popular pattern separation and pattern completion behavioural tasks in the literature (**Box S1**).

### Model output: “Same” or “different” decision

To model a delayed recognition task, such as the Occlusion at Retrieval Task, a decision on whether the recalled memory or model output pattern is the “same” or “different” to the original learned stimuli is required. One straightforward way to do this was to set a minimum accuracy level for the model to consider the pattern as being “same”. If the model output does not resemble the learned pattern, then the accuracy for that pattern would be low and would be considered as “different”. This, however, is not biological plausible as it requires the model to know the ground truth, i.e. correct answer, to compare with the model output pattern before making a decision.

As an alternative, we used the raw firing output of each neuron as a *certainty* measurement for whether that component of the pattern was ‘active’ or ‘inactive’. Based on the Hebbian learning principle, the stored synaptic values for active pattern components will be positive while the firing rate values for stored inactive pattern components will be negative. Therefore, the model will be more likely to recall a more positive firing rate neuron as ‘active’ and a more negative firing rate as ‘inactive’ whereas values in the middle may be considered either active or inactive by the model. This likelihood represents a *certainty* measure for the model. Since more positive and more negative values reflects greater certainty, we used the absolute value of each neuronal firing rate and applied a threshold to indicate whether the model is certain that the external input pattern matched a stored representation (Figure S6).

This threshold was set at the median firing rate to indicate an *unbiased prior* for choosing “same” or “different” and generally reflects having counterbalanced conditions within a psychological task. This threshold can be set differently according to task conditions. Furthermore, changing this threshold can reflect a bias which may arise with age or disease and this is explored later.

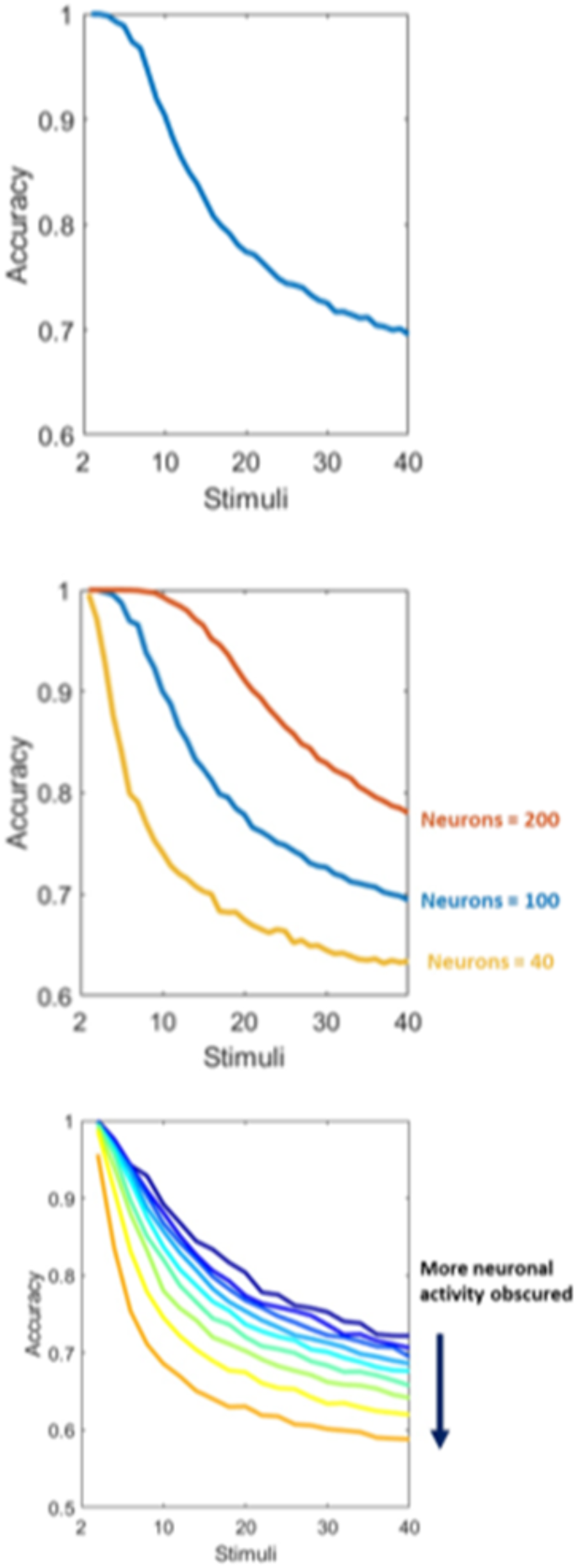

## Data availability

Data will be made available on reasonable request to the corresponding author Dr Xin You Tai

## Acknowledgments

This work was funded by the Wellcome Trust (Wellcome Trust PhD clinical fellowship to XYT and Wellcome Trust Principal Research Fellowship to MH) and MRC (Clinician Scientist Fellowship to S.M.) and by the Oxford NIHR Biomedical Research Centre.

## Author contributions

Design, analysis, manuscript writing, critical revisions (XYT); critical revisions of the manuscript (AS, MH); design, statistical analysis and critical revisions of the manuscript (SM)

