## Supplementary material for "Effects of focal epilepsy and aging on discrimination and reactivation of memories"

Table S1. Demographic data on cohort groups

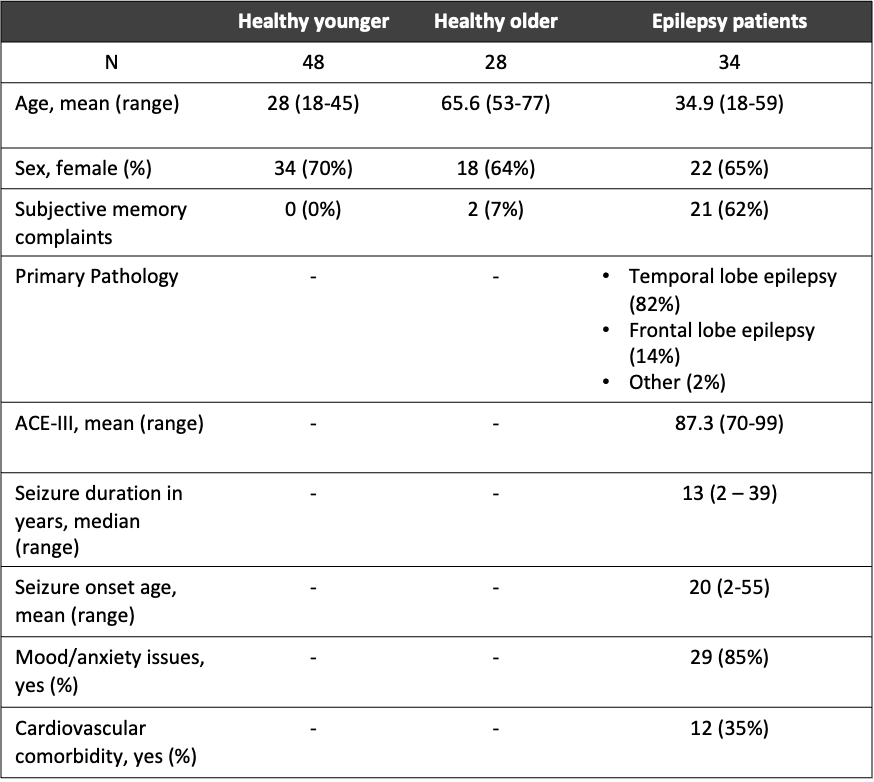

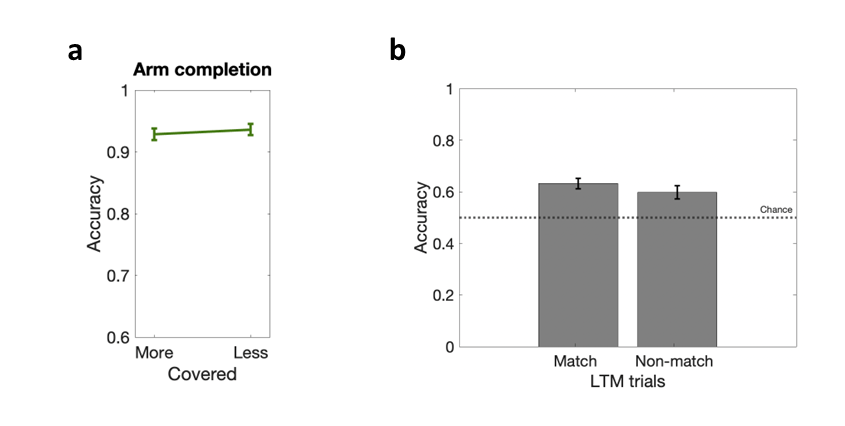

**Figure S1. Pinhole memory task active arm completion and long term memory results**

**a** Accuracy for active arm completion probes was higher with less covered probes. **b** For longer-term memory probes, accuracy was slightly higher for match probes compared with non-match probes. Error bars denote standard error of the group mean. PC – pattern completion, PS – pattern separation, LTM – longer-term memory.

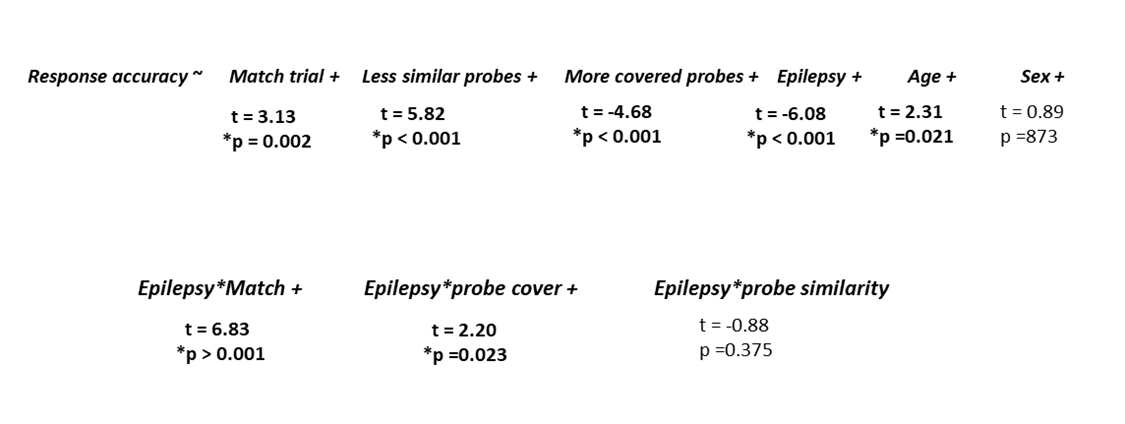

**Figure S2. Output of the Generalised Logistic Mixed effects model**

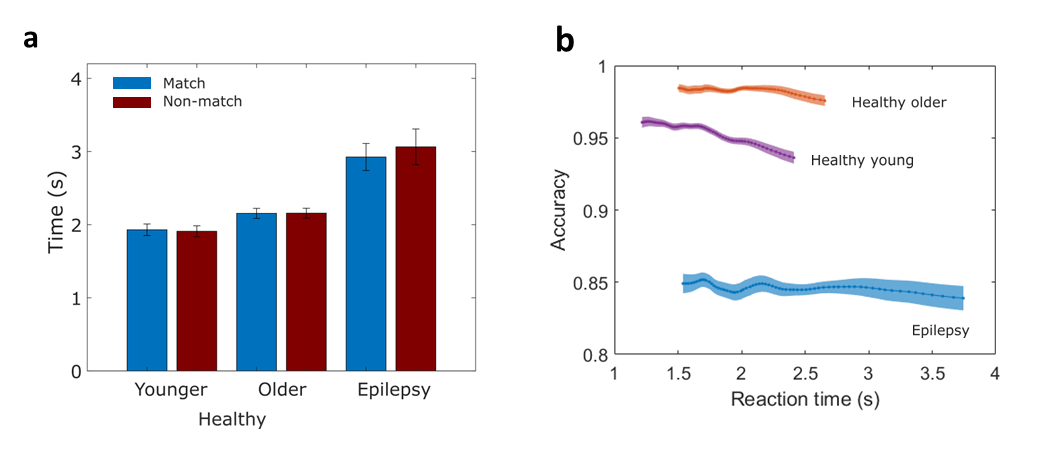

**Figure S3. People with epilepsy had the longest reaction time across participant groups**

**a** Within cohorts, there was no difference between match and non-match trial reaction times. Healthy young individuals had the fastest reaction times, followed by healthy older individuals and people with epilepsy. **b** Speed-accuracy trade-off visualised by plotting reaction time against accuracy.

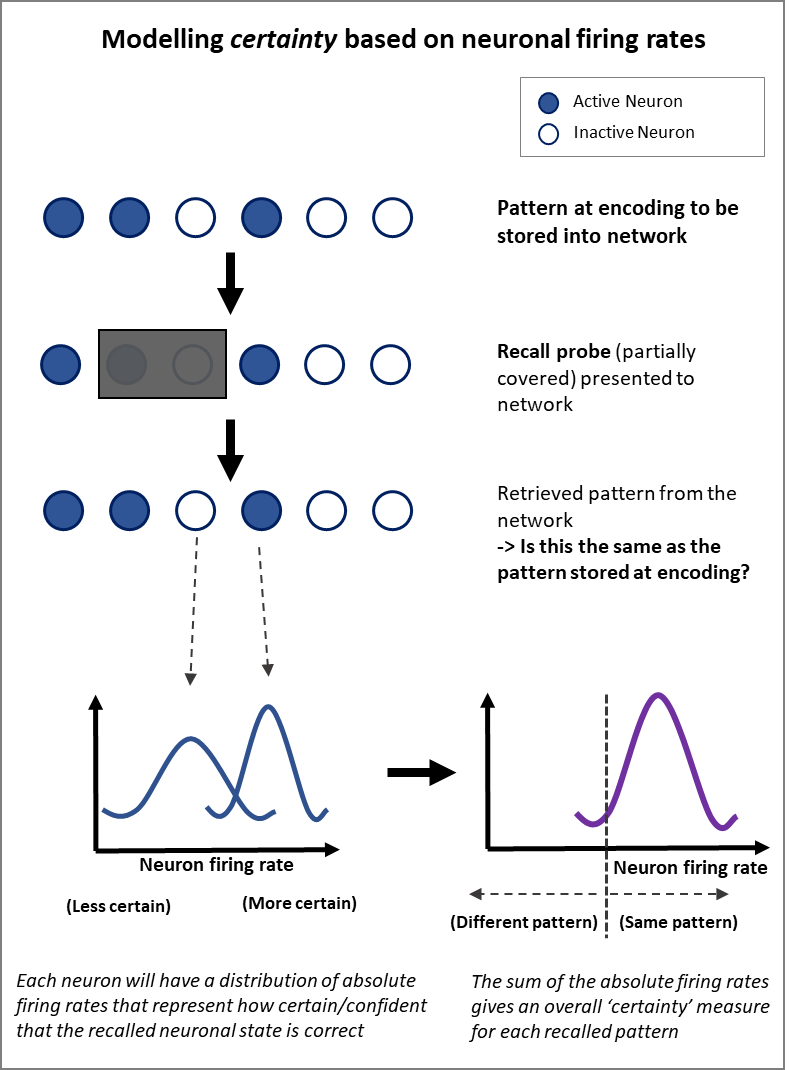

**Figure S4. “Same” or “different” decision based on model certainty**

Hopfield networks store patterns in synaptic weights at encoding. At a later stage, a recall probe will be presented to the model (this can be partially covered) and the network will retrieve a stored representation. Each neuron involved in this representation has a distribution of absolute firing rates that reflects the model certainty on whether that neuron was correctly recalled. These firing rates can be combined to give an overall certainty measure to decide whether the pattern was “same” or “different”.

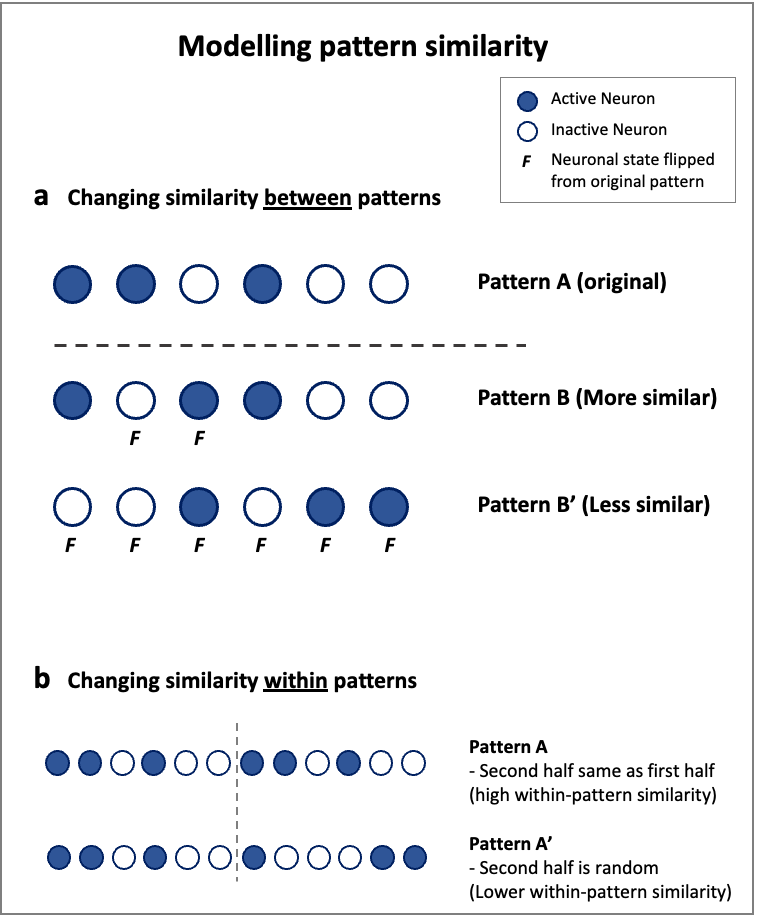

**Figure S5. Modelling different pattern similarity**

**a** Pattern A can be changed by flipping the state of one active and one inactive neuron to create a more similar pattern B (*between-pattern similarity*). Flipping the states of all the neurons will create a less similar pattern B’. **b** Modelling *within-pattern similarity* can be adjusted by having segments of repeat neuron activation patterns. Pattern A has high within-pattern similarity as the first six neuronal states are repeated whereas pattern A’ has lower within-pattern similarity with a random activation of the second six neurons.

**Box S1. Simulating results from popular pattern separation and pattern completion tasks from the literature**

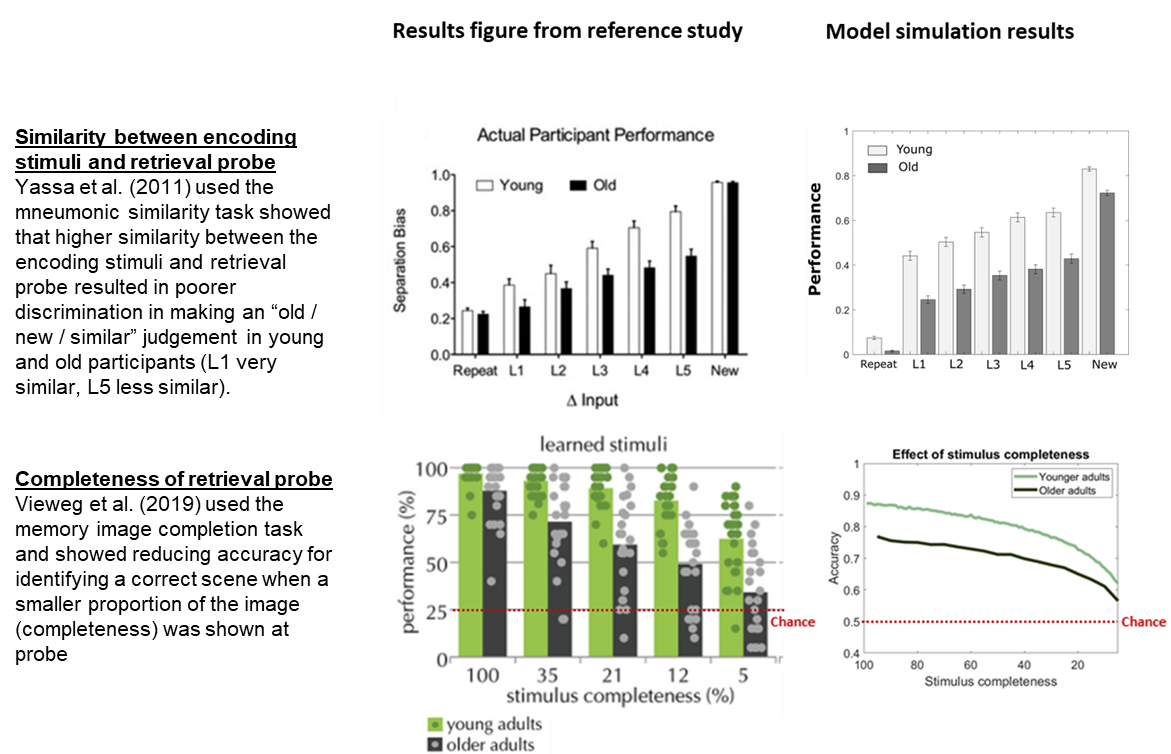
